# From language to pain: brain-behavior representations of hypnotic verbal suggestions for pain modulation

**DOI:** 10.64898/2026.07.22.740147

**Authors:** Dylan Sutterlin-Guindon, Marie–Eve Picard, Jen-I Chen, Mathieu Landry, Simona Brambati, David Ogez, Mathieu Piché, Pierre Rainville

## Abstract

Language can shape pain by conveying conceptual information about future somatosensory experiences. However, how verbal input is encoded in the brain and how these representations are later implemented during pain modulation remains unclear. We used a standardized hypnotic procedure with fMRI to test how verbal suggestions of hypoalgesia and hyperalgesia (vs neutral control) modulate pain. We quantified convergent neural responses across individuals during both suggestion encoding and pain processing, using inter-subject correlation and multivoxel pattern similarity approaches, respectively, and related these measures to individual differences in hypnotic suggestibility using representational similarity analysis. More suggestible individuals showed similar temporal dynamics in language-related regions during encoding and more similar hypoalgesia-specific patterns in the left parahippocampal gyrus (PHG) during pain. In contrast, lower suggestibility was associated with convergent responses in sensorimotor and monitoring systems across phases, demonstrating different neural implementation in individuals less successful in converting suggestions into hypoalgesia. These findings identify the PHG as a key region linking the effect of verbal suggestions to subsequent pain relief and show that hypnotic suggestibility structures how neural dynamics during language processing translate into altered pain experiences. This study offers a mechanistic lens for understanding how medical hypnosis and language-based therapies influence pain.

## Introduction

Language is a fundamental means of conveying contextual and pragmatic information about future experiences. Linguistic cues, including suggestions, instructions, and written information, can reliably modulate both the intensity and quality of pain. For example, placebo hypoalgesia can be induced by verbal statements such as “This medication should relieve your pain,” while a clinician’s wording may similarly shape patients’ pain experiences and recovery trajectories^1–4^. Such effects have been observed across multiple contexts, including placebo, learning, and hypnosis paradigms, demonstrating that language can influence pain and its neural processing^5–8^. Despite this evidence, most studies have focused on the effects of verbal information during pain or immediately before it, rather than on how such information is initially encoded. As a result, the neural processes linking language comprehension to pain modulation remain poorly understood.

Hypnosis provides an ideal experimental context to isolate the contribution of language to pain. Within hypnosis, verbal suggestions are instructions designed to alter aspects of cognition, affect, or perception^9^. These suggestions unfold dynamically over extended periods of time (i.e., minutes) and often convey meaning through metaphors that construct an alternative representation of bodily experience (e.g., “the arm feels light and numb, as if wrapped in a protective glove”). Such suggestions can produce pain relief (hypoalgesia) during surgery, improve chronic pain management, and consistently reduce pain in experimental settings^10–12^. Moreover, responsiveness to hypnotic suggestions varies substantially across individuals, with hypnotic suggestibility representing one of the strongest predictors of the hypoalgesic effects of verbal suggestions^12,13^. Hypnotic suggestibility is conceptualized as a trait reflecting the capacity to internalize and implement verbal instructions, making it a useful behavioral dimension for studying how language is transformed into perceptual change^14,15^. Although neuroimaging studies have consistently shown that hypoalgesic suggestions alter activity in pain-processing regions during painful stimulation^6,16–18^, the neural processes engaged while individuals listen to and encode verbal suggestions remain largely overlooked.

Suggestion encoding refers to neural processes involved in auditory and semantic processing, as well as the construction of conceptual representations from verbal input. These processes partly rest on regions of the language network, including the inferior frontal gyrus (IFG), superior temporal gyrus (STG), and middle temporal gyrus (MTG)^19^, which likely support conceptual representations of the suggested bodily state.

Once formed during suggestion encoding, these conceptual representations may influence pain via at least two mechanisms. First, expectancy-based mechanisms may translate conceptual representations into top-down predictions that influence subsequent pain processing^20,21^. Such effects are thought to involve prefrontal systems supporting attentional control and the affective regulation of instructed states^22,23^. Second, suggestion encoding may engage contextual learning mechanisms linking conceptual representations and expected sensory outcomes, supported by medial temporal lobe structures such as the hippocampus and parahippocampal gyrus (PHG)^24,25^. Consistent with this account, Desmarteaux et al.^26^ reported increased PHG activity during the encoding of pain modulatory suggestions, which predicted subsequent modulation of pain-related responses.

Together, these findings suggest that the encoding of suggestions involves language, expectancy and learning systems to produce subsequent pain modulation. However, the temporal order of these mechanisms, from suggestion encoding to downstream pain processing, and how this varies across individuals, remains unclear. Differential neural patterns associated with levels of hypnotic suggestibility may emerge during suggestion encoding, during pain processing, or across both phases, providing insight into how language-based pain modulation is actualized or counteracted.

In the present work, we conducted secondary analyses of fMRI data from Desmarteaux et al.^26^ to investigate how verbal suggestions are encoded in the brain and translated into pain modulation. We leveraged hypnotic suggestibility, assessed independently from the pain modulation protocol, as a behavioral dimension to characterize variability in these processes, reasoning that individuals who respond similarly to verbal suggestions may recruit similar neural mechanisms during the two key phases: 1) suggestion encoding and 2) pain processing. Participants (n = 23) listened to verbal suggestions designed to decrease or increase pain, followed by individually calibrated nociceptive stimulation (see Fig. 1).

**Figure 1.**
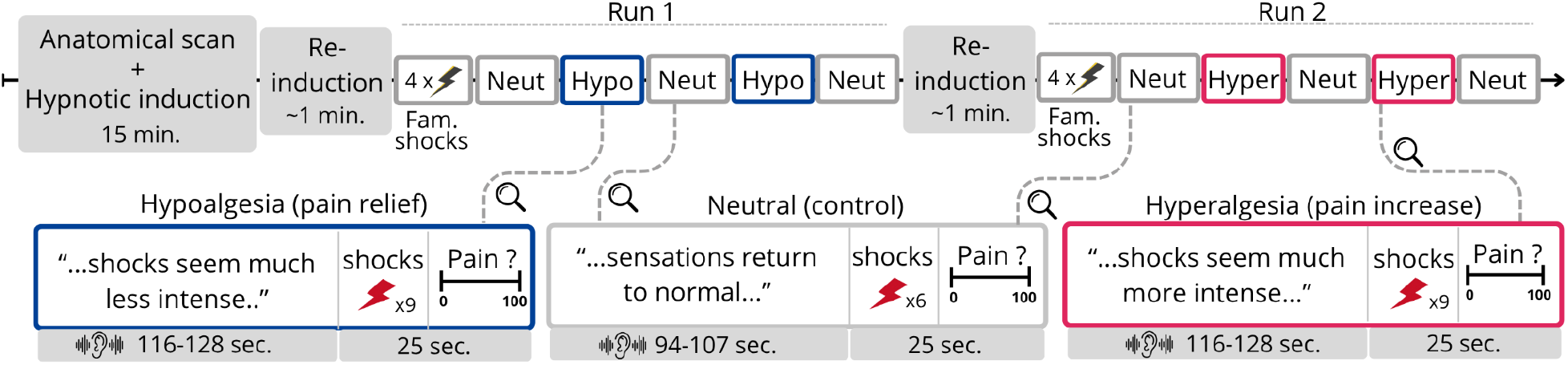
Experimental design of pain modulation induced by verbal suggestions. The study included twenty-three participants, who underwent two fMRI runs in a within-subject, counterbalanced design. A 15-minute standardized hypnotic induction procedure (based on the Stanford Hypnotic Susceptibility Scale form A; SHSS:A) was administered during the anatomical scan through MRI-compatible earphones. A 1-minute re-induction was administered prior to each run, followed by four familiarisation noxious stimuli (shocks). Painful electrical stimuli were individually calibrated and delivered to the right sural nerve. In each run, participants heard pre-recorded verbal suggestions designed to reduce pain (hypoalgesia, blue), increase pain (hyperalgesia, red), or feel pain normally (neutral control, gray). Each suggestion block was followed by a series of 6 or 9 shocks (ISI of 3, 6, or 9 sec.) and a pain rating using a 0-100 visual analog scale. Brain activity was recorded throughout and analyzed during two periods: the suggestion encoding phase (ear symbol), corresponding to the time participants listened to suggestions, and the pain induction phase (red lightning), corresponding to shock-evoked activity. Each run included a neutral suggestion condition repeated three times in alternation with the modulatory condition. These were labeled Neutral_HYPO_ or Neutral_HYPER_, depending on whether they occurred in the hypoalgesia or hyperalgesia run. Although identical in content, these neutral conditions were modeled separately to account for possible contextual influences on pain-related brain activity.

As shown in Fig. 2a, to characterize common and individual-specific neural processes across participants, we adopted an inter-subject framework that quantifies the degree to which behavioral and neural responses are similar, i.e., converge, across individuals^27^. Because verbal suggestions unfold as a dynamic stream of language, convergence in temporal patterns of brain activity across individuals should reflect common processes involved in transforming verbal input into conceptual representations of the suggested somatosensory state^28^.

**Figure 2.**
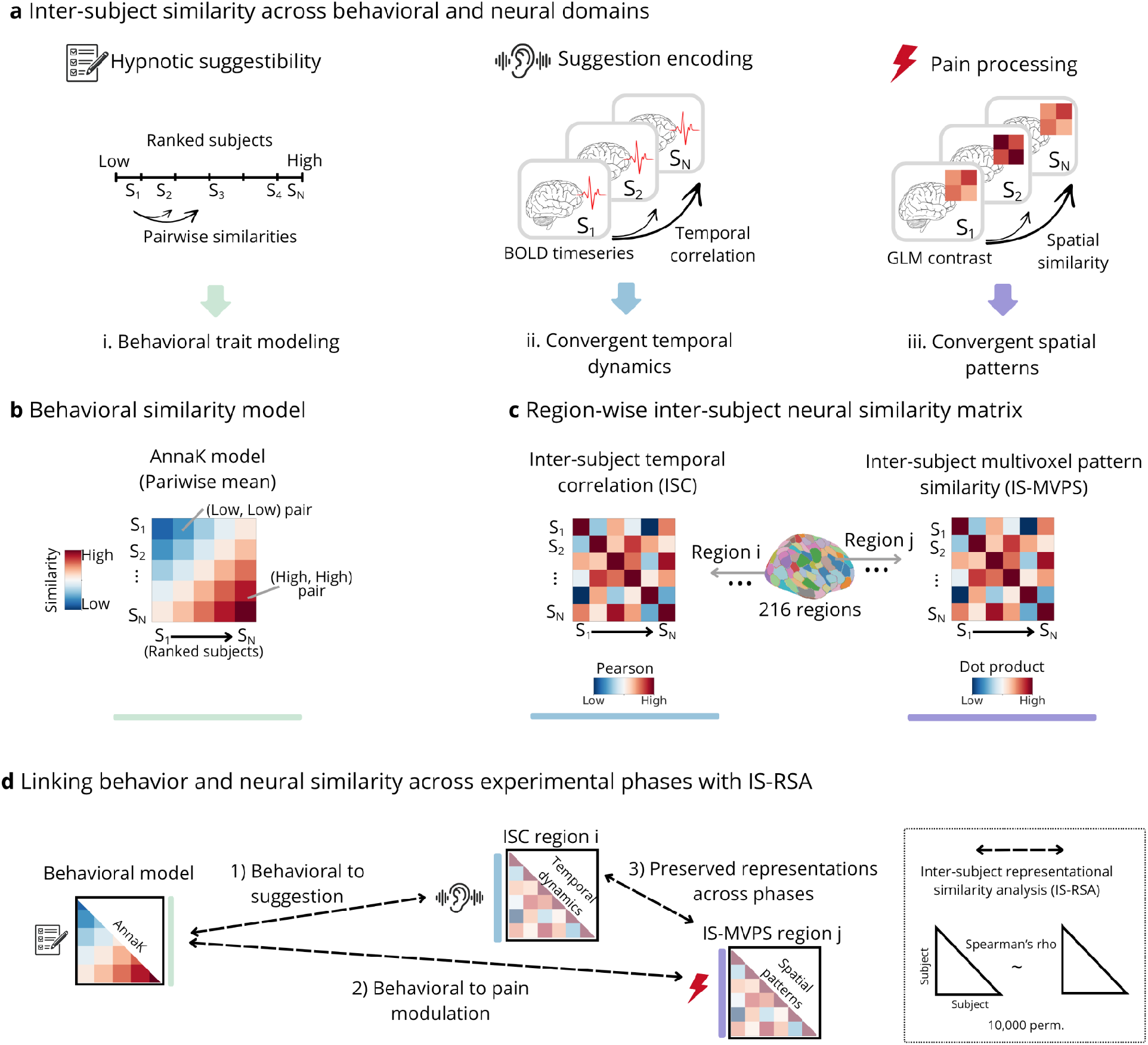
Inter-subject representational similarity framework. **(a)** We examined how hypnotic suggestibility relates to convergent brain responses across subjects during suggestion encoding (ear icon) and pain processing (lightning icon). We computed pairwise similarity between participants (n = 23 participants, for a total of 253 unique dyads) using hypnotic suggestibility scores, BOLD time series while participants listened to verbal suggestions, and multivoxel contrast maps during shock-evoked responses (e.g., Hypoalgesia vs Neutral_HYPO_). (i) Behavioral similarity was used to model the subsequent similarity in brain responses. (ii) Inter-subject correlations during suggestion encoding capture convergent temporal dynamics of brain activity. (iii) Inter-subject spatial correlations of pain-evoked patterns reflect convergent spatial patterns across individuals. **(b)** The hypnotic suggestibility model was built following the Anna Karenina principle (AnnaK;^27^). Similarity is computed from the mean of the suggestibility scores between each pair of individuals. Low-scoring dyads are expected to show more idiosyncratic/dissimilar brain patterns, while high-scoring dyads are expected to show similar brain patterns. **(c)** Inter-subject similarity matrices during suggestion encoding used Pearson-based inter-subject correlation (ISC), and during pain relied on inter-subject multivoxel pattern similarity (IS-MVPS), computed as the dot product between two individuals’ activity patterns in a given region. These matrices were computed independently for 216 regions. These approaches estimate the degree to which each brain region shows similar temporal or spatial patterns of brain activity between individuals. **(d)** Inter-subject representational similarity analysis (IS-RSA) was used to test whether patterns of behavioral similarity predicted patterns of neural similarity. Spearman rank correlations were computed between behavioral and neural similarity matrices to assess: (1) how suggestibility relates to ISC, (2) how suggestibility relates to IS-MVPS, and (3) functional representational link across phases between regions involved in suggestion encoding (e.g., region i found in 1) and regions involved in pain modulation (e.g., region j found in 2), revealing how neural representations of verbal suggestions are re-expressed during pain modulation as a function of hypnotic suggestibility. Statistical significance was assessed via 10,000 permutation tests.

During pain processing, we examined whether hypnotic suggestibility was associated with inter-subject convergence in spatial representations of pain modulation induced by suggestions. We hypothesized that if verbal suggestions engage common pain-regulatory mechanisms, individuals with similar responsiveness to suggestions would exhibit more similar multivariate representations during pain.

Finally, we tested whether convergence during suggestion encoding was associated with convergence during pain processing, reasoning that an inter-subject relationship between these stages would provide functional correlational evidence that the way verbal suggestions are encoded constrains how they are subsequently implemented into pain modulation.

We hypothesized that verbal suggestions would evoke convergent temporal responses within language and semantic networks during suggestion encoding, particularly among highly suggestible individuals. We further predicted that suggestibility-related convergence would emerge in spatial representations of pain modulation within brain systems implicated in expectancy and contextual learning. Finally, we expected individuals showing common encoding mechanisms during suggestion to also show similar expression of pain modulation, consistent with the idea that individual differences in the response to verbal suggestions partly arise from neurocognitive differences in how verbal suggestions are encoded.

## Results

### Behavioral results

At the group level, participants reported significantly less pain following hypoalgesic suggestions (Hypoalgesia > Neutral: *t*(22) = 3.72, *p* = .001) and more pain after hyperalgesia suggestions (Hyperalgesia > Neutral: *t*(22) = –2.75, *p* = .01). There was no difference in pain ratings between the two neutral conditions (Neutral_HYPO_vs. Neutral_HYPER_: *t*(22) = 0.22, *p* = .82), confirming the **s**pecificity of the suggestion effects (Fig. 3a).

**Figure 3.**
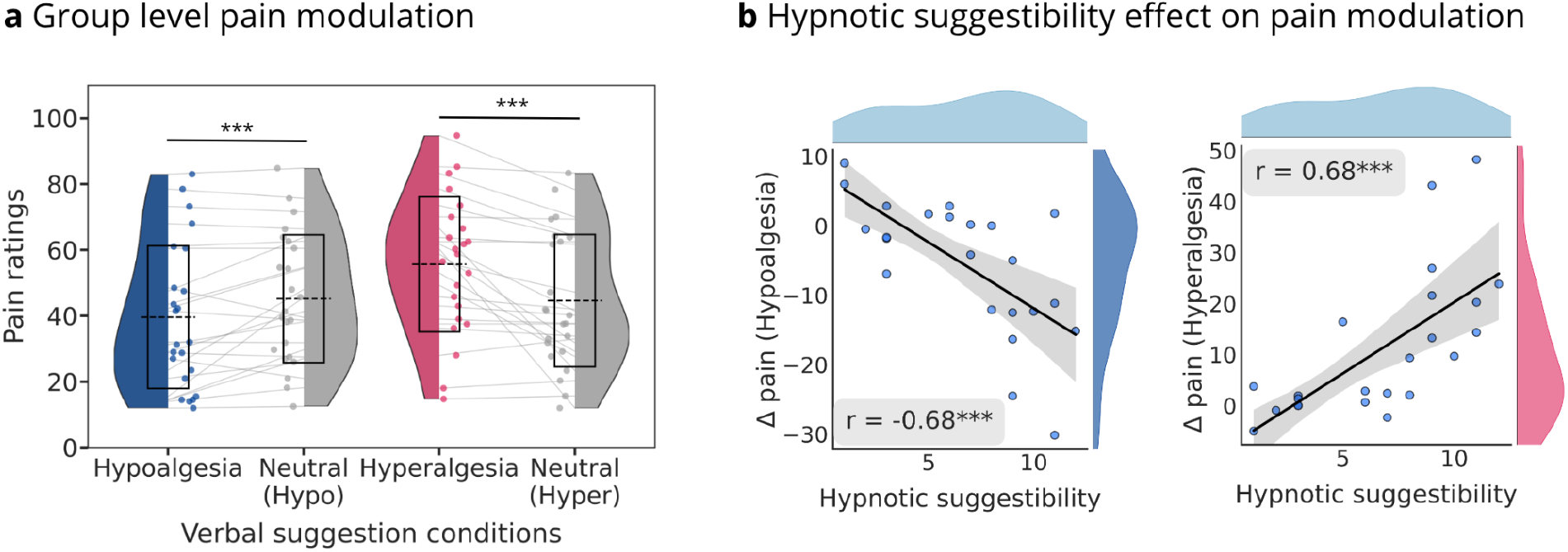
Mean pain ratings and individual differences related to hypnotic suggestibility. **(a)** Group-level pain modulation. Violin plots show pain ratings across suggestion conditions. Each dot represents an individual, and each line connects ratings from that individual across conditions. Boxes indicate the standard deviation, and the horizontal dashed line the mean. Pain ratings were significantly reduced during hypoalgesia compared to its matched neutral (Neutral_HYPO_), while pain ratings were significantly increased during hyperalgesia compared to its matched control (Neutral_HYPER_). **(b)** Individual differences in pain modulation as a function of hypnotic suggestibility. Scatter plots show correlations between hypnotic suggestibility (SHSS:A) and the magnitude of pain modulation for hypoalgesia (left) and hyperalgesia (right). Pain modulation for each subject (dot) was calculated as the difference between the mean pain ratings of the modulatory and the mean pain ratings of the neutral conditions (Hypo - Neutral; Hyper - Neutral). Higher suggestibility is associated with stronger pain reduction in the hypoalgesia condition (r = –0.675, p < .001) and stronger pain increase in the hyperalgesia condition (r = 0.681, p < .001). *** denotes p < .001.

Hypnotic suggestibility, as measured by the Stanford Hypnotic Susceptibility Scale version A (SHSS:A), significantly predicted how pain ratings were modulated by verbal suggestions. As depicted in Fig. 3b, higher SHSS scores were associated with greater pain reduction following hypoalgesic suggestions (*β* = –1.90, 95% CI [–2.85, –0.96], *t*(22) = –4.27, *p* < .0001) and stronger pain increase following hyperalgesic suggestions (*β* = 2.80, 95% CI [1.43, 4.16], *t*(22) = 4.27, *p* < .0001). Highly suggestible individuals showed the strongest pain modulation effects, while low suggestible individuals showed little or no pain modulation.

### Inter-subject correlation during suggestions encoding

To examine encoding processes while participants listened to verbal suggestions, we used intersubject correlation (ISC), which computes the correlation of a brain region’s BOLD time course between each pair of individuals. ISC reflects the degree to which temporal brain dynamics of a given region converge across subjects, revealing relevant systems engaged during suggestions^28^. Significant ISC was observed independently for each condition (Hypoalgesia, Hyperalgesia, and their matched control conditions Neutral_HYPO_ and Neutral_HYPER_; Fig. 4a), with the strongest effect observed in the bilateral (left > right) anterior superior temporal gyrus (aSTG; FDR-corrected, q < 0.05).

**Figure 4.**
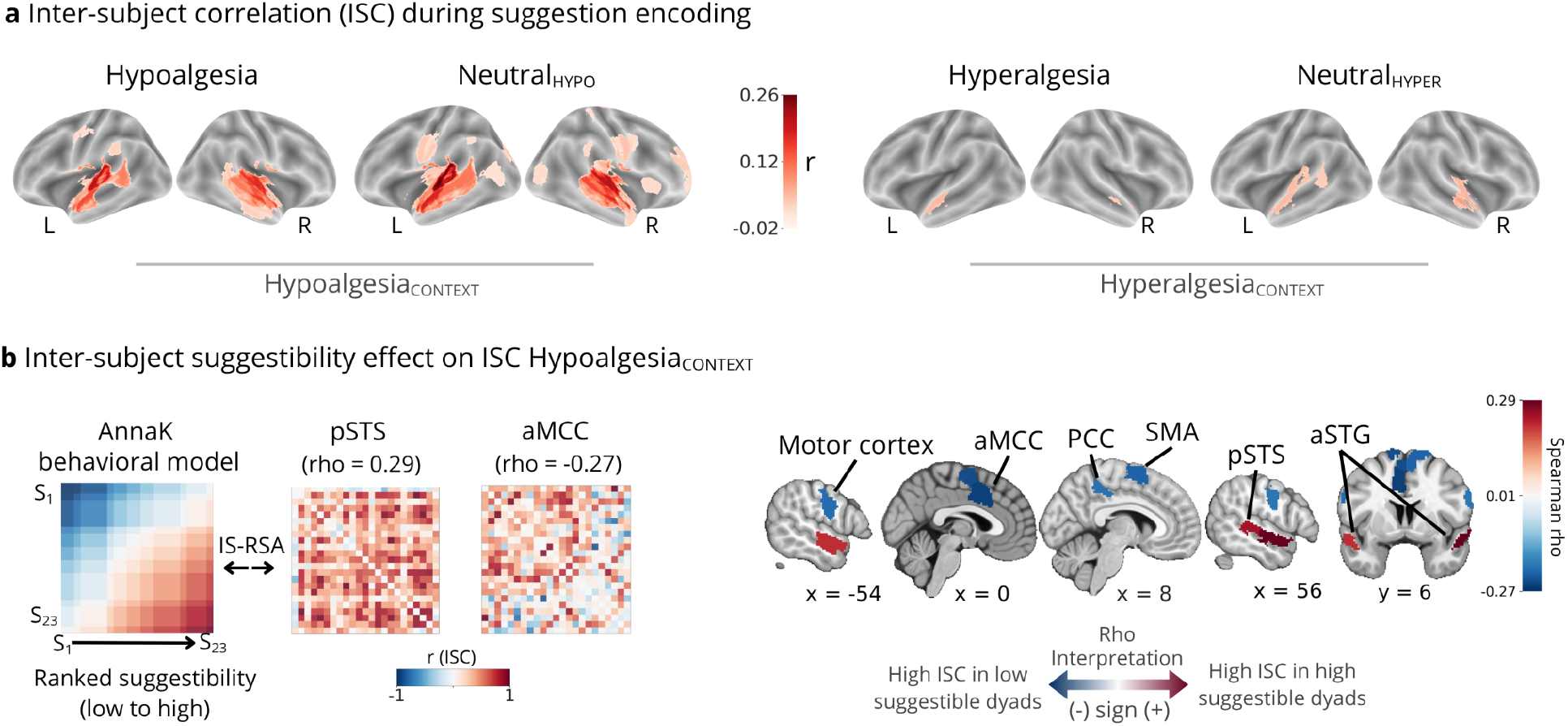
Inter-subject correlation during suggestion encoding as a function of hypnotic suggestibility. **(a)** Median ISC values across all subject pairs during listening, computed for each of 216 cortical parcels in the four conditions: Hypoalgesia, Neutral_HYPO_, Hyperalgesia, and Neutral_HYPER_. ISC values were tested against zero using permutation testing. Significant ISC was observed in bilateral temporal regions across all conditions (FDR-corrected, q < 0.05). A formal contrast revealed greater ISC during Hypoalgesia_CONTEXT_ compared to Hyperalgesia_CONTEXT_, supporting this condition grouping for subsequent analyses (see Supplementary Fig. S1). **(b)** Inter-subject representational similarity analysis (IS-RSA) tested the relationship between ISC matrices and the AnnaK behavioral model of hypnotic suggestibility during Hypoalgesia_CONTEXT_. Individuals with high suggestibility (red) showed greater ISC among them in the posterior superior temporal sulcus (pSTS) and anterior superior temporal gyrus (aSTG), while low suggestible individuals (blue) showed greater ISC among them in the motor cortex, supplementary motor area (SMA), posterior cingulate cortex (PCC), and anterior mid-cingulate gyrus (aMCC), revealing a specific contribution of these regions to one end of the suggestibility scale.

We next compared ISC across conditions on a region-by-region basis. Supplementary Fig. 1 and Supplementary Table S1 show that the Hypoalgesia_CONTEXT_ produced stronger ISC’s than the Hyperalgesia_CONTEXT_. Given that there were no significant differences between each modulatory suggestion and its matched neutral condition (e.g., Hypoalgesia > Neutral_HYPO_), we regrouped conditions according to these contextual effects, yielding: Hypoalgesia_CONTEXT_ (Hypoalgesia + Neutral_HYPO_) and Hyperalgesia_CONTEXT_ (Hyperalgesia + Neutral_HYPER_).

### Inter-subject representational similarity analysis (IS-RSA) during suggestion encoding

We next examined whether convergent temporal dynamics of brain activity during suggestion encoding were well explained by hypnotic suggestibility. To do so, we used inter-subject representational similarity analysis (IS-RSA) to characterize how ISC patterns were organized between dyads of participants in a given brain region. The Anna Karenina (AnnaK) model was used to test whether dyads of individuals with higher hypnotic suggestibility scores consistently display higher ISC (Fig. 2b-d). All reported results are FDR-corrected at q < 0.05.

Figure 4b shows that during suggestion encoding of the Hypoalgesia_CONTEXT_, ISC patterns of two broad sets of regions were well explained by the AnnaK hypnotic suggestibility model. First, we observed positive IS-RSA correlations in bilateral aSTG and the left posterior superior temporal sulcus (pSTS; peak ρ(251) = 0.29), indicating that individuals with higher hypnotic suggestibility encoded suggestions more similarly among them in these regions. Second, several regions associated with motor, somatosensory and monitoring functions showed negative IS-RSA correlations with the AnnaK model (peak ρ(251) = –0.27), including the anterior mid-cingulate cortex (aMCC), motor and somatosensory cortices, supplementary motor area (SMA), pre-SMA and the anterior portion of the posterior cingulate cortex (aPCC). These negative associations suggest that individuals with lower suggestibility exhibited a stronger ISC among them in these areas, while higher scorers showed more idiosyncratic responses.

During the Hyperalgesia_CONTEXT_, only the right supramarginal gyrus expressed more ISC among highly suggestible individuals (ρ(251) = 0.26, Supplementary Table S2). This aligns with the overall higher inter-subject variability observed in ISC in this context (Figure 4a).

To determine whether some convergent responses might be expressed across the entire suggestibility spectrum rather than preferentially among highly suggestible individuals, we conducted a complementary IS-RSA using a Euclidean-based similarity model. This analysis identified an additional effect in the left dlPFC during the encoding of Hypoalgesia_CONTEXT_ (Supplementary Results and Supplementary Fig. S2).

### IS-RSA of inter-subject multivoxel pattern similarity during pain modulation

We next examined whether inter-subject similarity in spatial activation patterns during pain processing was explained by hypnotic suggestibility. To address this, we applied IS-RSA to shock-evoked GLM contrast maps (Hypoalgesia > Neutral_HYPO_ and Hyperalgesia > Neutral_HYPER_), thereby reflecting brain response to pain modulation induced by verbal suggestions. For each brain region, we computed similarity between participants’ spatial patterns, yielding inter-subject multivoxel pattern correlation similarity (IS-MVPS) matrices. As in the prior analysis, we tested whether these neural similarity matrices were well explained by the AnnaK suggestibility model. Complementary analyses using the Euclidean similarity model are reported in the Supplementary Results and Supplementary Fig. S2. Significance is FDR-corrected at a rate q < 0.05, and full statistics for all models are reported in Supplementary Table S3.

For the inter-subject spatial similarity of the pain-evoked responses in Hypoalgesia vs Neutral_HYPO_, we found significant positive IS-RSA correlations between similarity in suggestibility and the left PHG (ρ(251) = 0.26), indicating that more suggestible pairs of individuals share more similar multivoxel hypoalgesia-related patterns (Figure 5a). A significant negative correlation was also found in the left SMA (ρ(251) = -0.26), indicating more convergent spatial responses among low suggestibility pairs of individuals in this region. Figure 5b illustrates the AnnaK structure of the left PHG, where the direction and magnitude of multivoxel patterns within this region tend to converge more among pairs of individuals with higher suggestibility scores.

**Figure 5.**
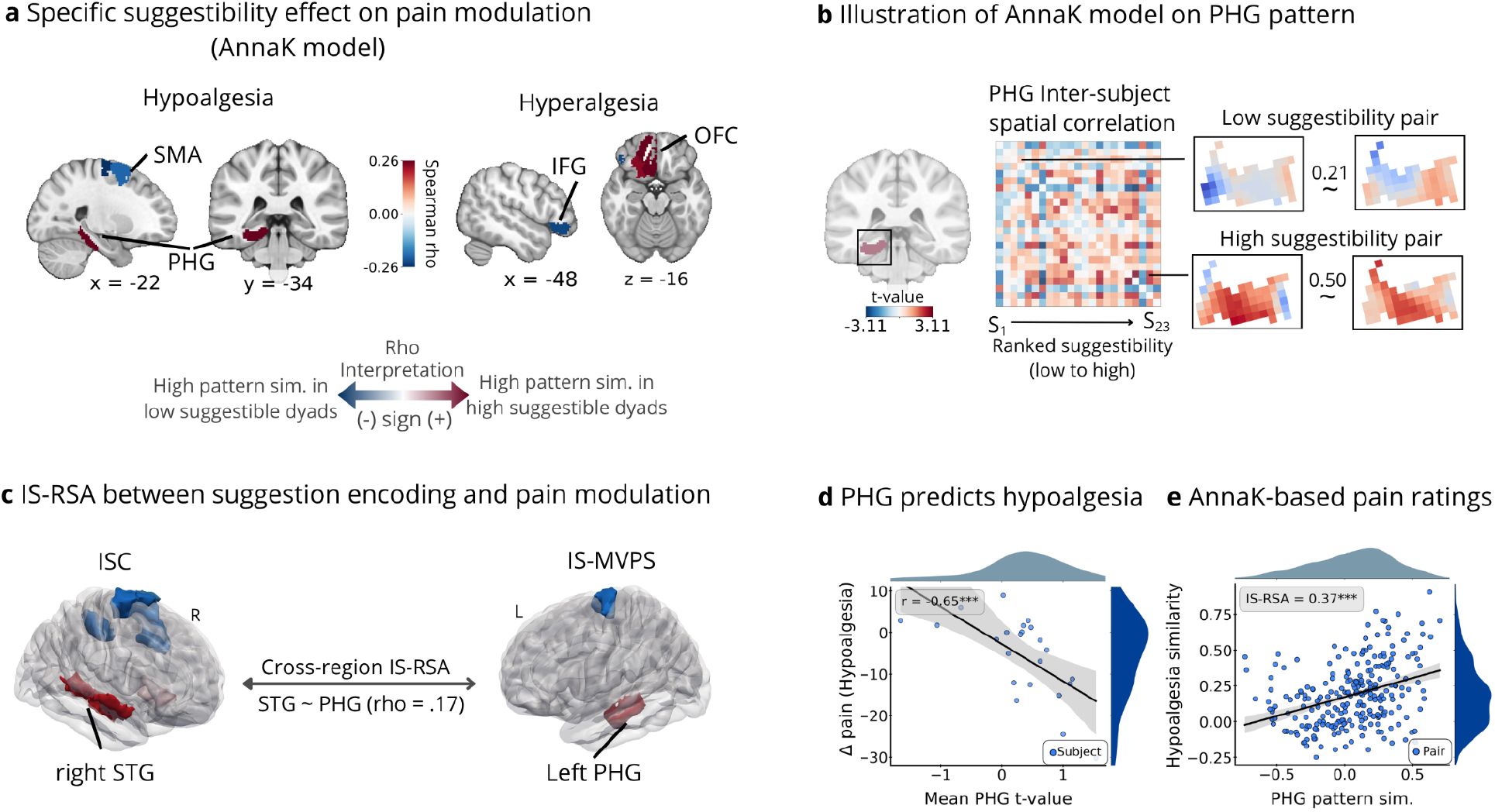
Inter-subject multivoxel pattern similarity (IS-MVPS) during pain modulation. **(a)** Specific contribution of hypnotic suggestibility scores (AnnaK model) to IS-MVPS during hypoalgesia and hyperalgesia. IS-MVPS was computed from shock-evoked contrast maps (Hypoalgesia > Neutral_HYPO_ and Hyperalgesia > Neutral_HYPER_). Regions shown in red indicate that highly suggestible individuals shared more similar multivoxel modulation patterns among them, while regions in blue indicate greater pattern similarity among low suggestible individuals. **(b)** The parahippocampal gyrus (PHG) is highlighted as a region showing suggestibility-specific effects during hypoalgesic modulation. For illustration, PHG multivoxel hypoalgesia patterns are shown for a low suggestibility pair (similarity = 0.21) and a high suggestibility pair (similarity = 0.50), showing a general trend of increased representational overlap in high responders. **(c)** Preserved inter-subject structure from suggestion encoding to pain modulation. IS-RSA between inter-subject correlation (ISC) patterns and the IS-MVPS in regions having previously shown a positive AnnaK structure. The right anterior and posterior STG showed a positive fit with the left PHG, indicating that individuals with similar temporal dynamics during suggestion in the right STG tend to show a similar spatial pattern in the left PHG during hypoalgesia. **(d)** Relationship between PHG spatial patterns and behavioral hypoalgesia. A subject-wise analysis shows that mean PHG t-values during hypoalgesia are negatively correlated with behavioral hypoalgesia (Δ: Hypoalgesia - Neutral_HYPO_ ratings) (r = 0.65). **(e)** AnnaK-based modeling of behavioral pain modulation. IS-RSA between this AnnaK model and IS-MVPS in the left PHG shows that pairs of individuals with more similar behavioral hypoalgesic modulation also showed more similar PHG spatial patterns during hypoalgesia (rho = 0.37). Supplementary motor area (SMA); parahippocampal gyrus (PHG); Inferior frontal gyrus (IFG); Orbitofrontal cortex (OFC); Superior temporal gyrus (STG).

For the inter-subject spatial similarity of the pain-evoked responses in Hyperalgesia vs Neutral_HYPER_, as shown in Supplementary Table S3, we found positive IS-RSA effects in two regions of the orbitofrontal cortex (OFC; peak ρ(251) = 0.26; Fig. 5a) and in the right motor area (ρ(251) = 0.22). This indicates greater hyperalgesic pattern similarities among more suggestible participants in those areas. In contrast, a negative correlation was found in the left inferior frontal gyrus (IFG; ρ(251) = -0.23; Fig. 5a), indicating more similar spatial patterns in less suggestible participants.

### Cross-region IS-RSA linking suggestion encoding to pain modulation

The previous analyses showed that individual differences in suggestibility were reflected in ISC during suggestion encoding and in spatial patterns during pain modulation, across partially distinct sets of regions. We therefore tested whether convergent inter-subject neural patterns established during suggestion encoding were preserved in regions implicated in hypoalgesia using a cross-region IS-RSA framework (Fig. 2d). Seed regions were defined based on significant IS-RSA effects during suggestion encoding, and target regions based on significant IS-RSA effects during pain modulation (see Supplementary Table S4).

Figure 5c illustrates how ISC in the right aSTG and right pSTG during Hypoalgesia_CONTEXT_ suggestions correlated with spatial pattern similarity in the left PHG during hypoalgesic pain (aSTG-PHG: ρ(251) = 0.14, p = 0.01; pSTG-PHG: ρ(251) = 0.17, p = 0.003; FDR-corrected at q < 0.05; see Supplementary Table S5). Both STG and PHG had independently shown a positive fit with the behavioral AnnaK suggestibility model in their respective analyses (ISC and IS-MVPS), indicating that individuals who encoded hypoalgesic suggestions more similarly in these temporal regions also exhibited more similar spatial patterns in the left PHG during pain modulation, especially among individuals with higher suggestibility.

### Extension of IS-RSA to behavioral changes in pain ratings

Having used hypnotic suggestibility as a behavioral dimension to identify brain regions preferentially showing convergent multivoxel patterns across subjects during pain modulation, we confirmed that these regions were also related to behavioral modulation of pain ratings. We first tested whether left PHG activity was directly associated with behavioral pain modulation. Figure 5d shows that mean activation in the left PHG during Hypoalgesia versus Neutral_HYPO_ was negatively correlated with the magnitude of hypoalgesia (r(22) = -0.65, p < 0.001), indicating that individuals showing stronger PHG activation during hypoalgesia experienced greater reductions in reported pain compared to the control condition.

We then examined whether multivoxel pattern similarity in the left PHG captured inter-individual differences in behavioral pain modulation. Using an IS-RSA approach, we constructed an AnnaK similarity model based on changes in pain ratings (Hypoalgesia - Neutral_HYPO_) and tested its overlap with inter-subject similarity in PHG activation patterns. Figure 5e shows that PHG pattern similarity was significantly correlated with this behavioral model (ρ(251) = 0.37; Supplementary Table S6), indicating that individuals with more similar PHG representations also showed more similar magnitudes of pain relief.

## Discussion

Language can convey conceptual representations that shape pain, yet the neural processes linking the encoding of hypnotic verbal suggestions to subsequent pain modulation remain poorly understood. Using hypnotic suggestibility as a behavioral dimension indexing how individuals translate verbal suggestions into perceptual changes, we examined inter-subject convergence in neural representations during both suggestion encoding and pain processing. Our results show that individuals who exhibited more similar neural responses while listening to suggestions also tended to exhibit more similar neural and behavioral patterns of pain modulation, providing evidence for a functional link between language processing during suggestion encoding and downstream pain-regulatory mechanisms.

During suggestion encoding, individuals with higher hypnotic suggestibility expressed convergent neural dynamics within language-related regions, while they exhibited convergent spatial representations of hypoalgesia within the left parahippocampal gyrus (PHG), suggesting that successful pain relief by verbal suggestion partly depends on how conceptual representations are constructed during the listening phase. In contrast, among individuals with lower suggestibility, we observed convergent neural responses in regions involved in bodily monitoring, motor control, and action-related processing during both encoding and pain. The neural processes at the encoding stage, organized by hypnotic suggestibility, appear to influence the implementation of verbal suggestions into pain modulation across individuals. More broadly, these findings provide a framework for understanding how psychological traits shape the transformation of language and hypnosis into altered subjective experience across distinct stages of neural processing.

More suggestible individuals exhibited greater inter-subject correlations in the bilateral anterior superior temporal gyrus (aSTG) and right posterior superior temporal gyrus (pSTG) during the Hypoalgesia_CONTEXT_, indicating more convergent neural responses while processing the suggestions. These regions are part of a high-level language and semantic network involved in extracting meaning from verbal input and linking linguistic content to conceptual knowledge^19,29^. Although these effects cannot be attributed specifically to the hypoalgesic content of the suggestions, because of the inclusion of the control condition, they indicate that hypnotic suggestibility shapes how verbal information delivered in a hypnotic context is encoded. More specifically, the AnnaK model revealed that the geometry of these inter-subject similarity patterns was asymmetrical across the suggestibility spectrum (but see supplementary results and discussion for the Euclidean-based results). Individuals with higher suggestibility showed increasingly similar temporal dynamics among them in higher-order language regions, whereas lower-suggestibility individuals exhibited more heterogeneous responses. This convergence may therefore reflect common cognitive operations engaged while constructing conceptual representations of the suggested bodily state^28^.

Importantly, convergence during suggestion encoding was associated with convergence in pain-related representations within the left PHG. Individuals who exhibited more similar inter-subject correlations in the right anterior and posterior STG also showed more similar multivoxel pain-related patterns in the PHG during hypoalgesia. This relationship suggests that the way verbal suggestions are encoded constrains how they are subsequently implemented during pain processing. Structural and functional connections between temporal language regions and the PHG^30^ provide a plausible pathway through which conceptual representations formed during listening may influence downstream pain-modulatory systems.

Notably, both the encoding effects and their relationship to left PHG representations were strongest in the right hemisphere. Right temporal regions, especially the aSTG, have been implicated in processing linguistic information over extended contextual windows and integrating semantic information across longer timescales than homologue left regions, particularly during naturalistic language comprehension^31^. These regions also contribute to pragmatic aspects of communication^19,32^, which are central to hypnotic suggestions. This lateralization therefore suggests that successful suggestion encoding may depend on context-sensitive and pragmatic aspects of language processing rather than solely semantic components.

During pain processing, the left parahippocampal gyrus (PHG) emerged as a key region expressing hypoalgesic representations. Participants showing the strongest pain relief from hypoalgesic suggestions showed more similar multivoxel activity patterns in the left PHG, whereas lower responders showed more idiosyncratic patterns. In addition, the level of PHG activation during hypoalgesia predicted the magnitude of pain relief at the individual level. These effects emerged contralaterally to the nociceptive stimulation, and while controlling for activity during Neutral_HYPO_ trials, reinforcing the specific involvement of the left PHG in pain relief. The PHG has been associated with reduced pain perception at a given stimulus intensity^33^, is a core component of the stimulus intensity independent pain signature, a predictive model of endogenous pain modulation^34^, and co-varies with regions involved in descending pain control, such as the periaqueductal gray^35^. Its recruitment during placebo analgesia^36^ further supports its role in verbally induced hypoalgesia and expectancy-based processes^20^. Together, these observations suggest that the left PHG is a key region for enacting pain inhibition induced by hypoalgesic verbal suggestions.

The present findings extend previous analyses of the same dataset^26^, which identified increased left PHG activation during the encoding of pain-modulatory versus neutral suggestions, and subsequently predicted pain-evoked patterns in nociceptive regions. While that work demonstrated PHG involvement during suggestion encoding, the present results show that PHG representations during pain processing are themselves organized by hypnotic suggestibility, relate to encoding processes, and to behavioral pain modulation. These results suggest that the PHG is consistently engaged in representing verbally induced contextual information, and that its role extends from encoding suggested states to their implementation during pain modulation.

Our study supports a conceptual and associative account of pain modulation induced by verbal suggestion. The PHG is also central to contextual memory, associative integration, and the encoding of relational contingencies^24,37^. It supports the retrieval of verbally learned contextual information^38^, and links semantic content to self-relevant contexts^39^. Within predictive-processing frameworks^40,41^, verbal suggestions may establish conceptual representations of future bodily states during listening. In more suggestible individuals, these representations may be encoded more consistently within language networks and subsequently reactivated within contextual-associative systems such as the PHG during pain. This activation during pain may reflect the reinstatement of verbally suggested context (e.g., “your foot feels protected”) and its integration with incoming sensory input, thereby engaging top-down modulation via connections with the endogenous pain regulation systems.

One possibility is that verbal suggestions operate through a form of one-shot learning, whereby a single episode of linguistic input establishes predictive links between contextual representations and nociceptive processing^42^. This interpretation aligns with evidence that verbal instructions alone can modify pain perception without reinforcement^1,8^. More broadly, the findings support theories proposing that language can convey sensori-affective meaning through conceptual associations, allowing verbal information to shape future somatosensory experience^43–45^.

In contrast to highly suggestible individuals, low responders, who showed weaker behavioral pain modulation, exhibited convergent neural patterns during suggestion encoding in distinct sets of regions. During Hypoalgesia_CONTEXT_, they showed convergent temporal dynamics in sensorimotor and monitoring-related regions, including the aMCC, sensorimotor cortices, SMA, and pre-SMA. These regions are commonly associated with action preparation, intentional control, and self-monitoring^46,47^.

This pattern suggests that low responders may engage verbal suggestions through more controlled processing pathways. The aMCC, often co-activated with SMA and pre-SMA, has been implicated in volitional action, self-monitoring ^48^, and metacognitive judgment^49^. Correlated patterns of activity in these regions may therefore reflect a common tendency to engage with suggestions through monitoring or internally guided control processes, rather than through the more convergent automatic processing typically observed in highly suggestible individuals^50^. This interpretation is consistent with prior work linking suggestibility to differences in attentional and monitoring processes^51^.

In parallel, low responders showed more heterogeneous inter-subject correlations in temporal language regions, suggesting more idiosyncratic processing of the semantic content of suggestions. This variability at the encoding stage may contribute to weaker or less consistent downstream effects on pain modulation. During pain, low responders also exhibited convergent spatial patterns in the left SMA, indicating a reliance on motor-preparatory systems during pain rather than associative representations in regions such as the PHG.

Together, these findings advance the idea that if relevant conceptual representations are not established during suggestion encoding, downstream pain modulation relies more heavily on monitoring and preparatory systems. This alternative processing mode may limit the extent to which verbal suggestions can engage higher-order associative mechanisms to shape pain, favouring instead bottom-up nociceptive signals.

Our results also identified regions important for hyperalgesia. During pain processing, orbitofrontal cortex (OFC) regions showed inter-subject spatial pattern similarity among more suggestible individuals, while showing heterogeneous patterns in lower responders. The OFC is involved in integrating affective and contextual information and encoding the subjective value of pain^52^. It is also engaged during cognitive reappraisal and expectation-based modulation of affective states, and is functionally connected to descending pain-control systems^53^. In this context, convergent OFC representations may reflect a common value-based reappraisal process during hyperalgesic suggestions, supporting the reinterpretation of nociceptive input in a way that amplifies the pain experience. Along with the OFC regions, we found that the anterior midcingulate cortex (aMCC) also contributed to how hyperalgesia was expressed across individuals, but through a distinct inter-subject representational structure (See supplementary results and discussion). Altogether, these results suggest that pain amplification can be achieved through distributed affective systems in which each region has its own interaction pattern with hypnotic suggestibility levels.

This study has implications for clinical settings. Verbal suggestions are central to many therapeutic interventions, including medical hypnosis and psychotherapy. The present findings provide a mechanistic framework in which the way verbal suggestions are encoded constrains how they are later implemented in cognitive-affective systems. In particular, our findings indicate that variability in treatment response may arise from differences in how patients initially process therapeutic language and that this variability can be indexed by an objective phenotypical measure.

These mechanisms are especially relevant in acute and chronic pain, where expectations and learned associations shape symptom expression^54^. Rather than acting solely at the level of downstream regulation, language-based interventions may operate by establishing new contextual and associative representations that are later recruited during pain processing. This perspective aligns with evidence that verbal instructions can modify pain-related beliefs and reduce maladaptive responses such as fear and avoidance^55^. Language-based therapies may help restructure maladaptive schemas by targeting the PHG and by fostering new contextual associations.

We also find that individual differences, such as hypnotic suggestibility, influence how verbal suggestions are encoded and translated into changes in pain perception. While this trait is a strong predictor of hypnotic analgesia in clinical and experimental settings, other individual characteristics, such as attention efficiency, mental imagery ability, or phenomenological measures of hypnotic experiences, may also contribute to the inter-individual variance in how verbal suggestions are encoded and translated into changes in experience^51,56^. Future work should examine how these factors shape the consistency of neural responses during encoding and their downstream effects on pain. Investigation of these additional variables could unveil useful clinical predictors of treatment response, leading towards the development of personalized understandings of hypnotic analgesia.

The present study leveraged naturalistic stimuli methodologies to examine how extended language input is encoded and later expressed during pain. While this approach captures ecologically valid aspects of a therapeutic intervention, it relies on a limited set of suggestion types presented identically across participants. Future studies could systematically vary the structure and content of verbal suggestions, such as purely literal versus metaphorical, to determine how these features shape the formation of conceptual representations during hypnosis. Extending this approach to other affective or interoceptive domains would further clarify the generality of the observed functional trajectories of the process.

More broadly, the present findings motivate a shift from group-average approaches toward models that explicitly capture inter-individual variability in how dynamic cognitive processes are implemented across distinct systems and processes. While the current sample size is admittedly modest, the use of inter-subject representational analyses provides a framework for leveraging pairwise structure (n=253) to study such variability across different brain states and experimental phases. Replication in larger samples and across independent datasets, including clinical samples, will be essential to establish the robustness of these effects and to determine how generalizable these associative conceptual processes influence pain modulatory mechanisms.

### Conclusion

This study identifies a link between the brain encoding of verbal suggestions and their later expression during pain. Across individuals, similarities in brain responses while listening to suggestions were associated with similarities in pain-related representations, particularly during hypoalgesia. This relationship was structured by hypnotic suggestibility, indicating that individual differences organize how consistently suggestions are encoded and implemented in the brain. The parahippocampal gyrus emerged as a key site of hypoalgesia and was related to behavioral pain relief, likely supporting contextual associations between verbally induced meaning and pain modulation. In parallel, distinct encoding trajectories in low suggestibility individuals point to an alternative pathway relying more on monitoring and sensorimotor processes. These findings show that the way verbal suggestions are initially processed is associated with the magnitude of subsequent pain modulation. Together, this study provides an account of how language-based information can be transformed into changes in affective experience.

## Methods

### Participants

The sample consisted of 24 healthy participants included in the study of Desmarteaux et al.^26^ One participant was excluded in this reanalysis because of missing time series data, making inter-subject correlations (see below) invalid. The final data set included 23 individuals (12 females, 11 males; mean age = 26.8 years, SD = ±1.1). All procedures in the original study and in the secondary use of the data reported here were approved by our local ethics committee and complied with the Declaration of Helsinki. All participants provided informed consent for participating to the study. At the time of data collection, the consent form did not include authorization for the sharing of individual data outside of the research group.

Participants took part in two visits, one to assess hypnotic suggestibility and to get familiar with the noxious stimulation procedure, and another for the fMRI experiment. Hypnotic suggestibility was assessed using the French version of the Stanford Hypnotic Susceptibility Scale, Form A (SHSS:A)^57^. This individually administered procedure takes approximately 30-45 minutes and provides a standardized measure of responsiveness to hypnotic suggestions.

### Overview of the brain imaging procedure

The scanning procedure is illustrated in Fig. 1 and was acquired with a 3T MRI. The induction of hypnosis was adapted from the procedure included in the SHSS:A and pre-recorded to be delivered via earphones during the structural scan (i.e., high-resolution T1 image; see imaging acquisition details in^26^). This was followed by two functional scanning runs (T2* EPI, BOLD), each including two blocks of hypoalgesia or hyperalgesia alternating with three blocks of a control condition with neutral suggestions (order of hypoalgesia and hyperalgesia runs counterbalanced between participants). Within each block, auditory verbal suggestions were delivered, followed by a series of 30-ms transcutaneous noxious electrical stimulation applied to the right sural nerve (n=36 pain stimuli per run, allocated equally to the pain modulation and the neutral condition).

### Hypnotic verbal suggestion

During the fMRI runs, participants listened to three types of pre-recorded verbal suggestions: suggestions to decrease pain (Hypoalgesia), to increase pain (Hyperalgesia), and to maintain normal pain perception (Neutral). In the Hypoalgesia condition, participants were instructed to imagine their foot transforming into rubber, a material known for its insulating properties. For example: “Your skin becomes numb and you will hardly feel any stimulation. It’s like a layer of rubber between your skin and the stimuli.” In contrast, the Hyperalgesia condition invited participants to imagine their foot becoming metallic, emphasizing conductivity and heightened sensation: “Your skin is becoming very sensitive and you will feel more and more stimulation. It’s as if a layer of metal were placed between your skin and the stimuli.” In the Neutral condition, suggestions emphasized a return to normal bodily state and sensation: “Your ankle is no longer numb and your foot is back to its normal constitution of bone, muscle and flesh.”

The hypoalgesic and hyperalgesic suggestions were carefully designed to maintain comparable sentence structure, tone, and affective engagement. Suggestion blocks varied slightly in duration: with three blocks consisting of the neutral suggestions ranging from 94 to 107 seconds (Neutral total per run = 300 sec), and two blocks consisting of modulatory suggestions (Hypoalgesia and Hyperalgesia) ranging from 116 to 128 seconds (Hypoalgesia total = 247 sec; Hyperagesia total = 239 sec). Although the neutral suggestions administered in the hypoalgesia and hyperalgesia runs were identical, they were treated separately to account for the context embedding of each run (i.e. Neutral*_HYPO_*, and Neutral_HYPER_; also see analysis below). Full verbatim transcripts of the suggestions are provided in the Supplementary Materials in^26^.

### Painful stimulations and behavioral ratings

After the delivery of verbal suggestions in each condition, participants received a series of 6 to 9 transcutaneous electrical stimulations on the left sural nerve with the inter-stimulus interval (ISI) varying pseudorandomly between 6, 9 or 12 sec. Each stimulus was individually calibrated to elicit a reliable nociceptive reflex and moderate pain intensity. Pain intensity and unpleasantness were rated on a visual analog scale (0-100).

### Analyses

fMRI data preprocessing was performed on SPM8 (Statistical Parametric Mapping, Version 8; Wellcome Department of Imaging Neuroscience, London, UK). Preprocessing steps included slice timing correction, motion correction and re-slicing with fourth-degree B-spline interpolation. Images were spatially normalized to MNI space using structural images for parameter estimation of the unified-segmentation method. Spatial smoothing was applied to the functional time series using a 6-mm isotropic full width half maximum Gaussian kernel. Finally, a high-pass temporal filter at 428 s and a correction for auto-correlation (AR1) were applied to the time series data.

Data analyses were implemented through custom Python 3.11 scripts, openly available at https://github.com/dylansutterlin/ISC_hypnotic_suggestions. Main software packages used to perform the analyses include BrainIAK and Nilearn.

### Hypnotic suggestibility and Pain ratings

For each pain trial (i.e., the series of painful stimuli following a given suggestion), we computed the mean of the intensity and unpleasantness ratings, as these two measures were highly correlated across subjects (r’s = 0.76–0.95). Paired t-tests were used to evaluate differences in pain ratings between each modulatory suggestion (Hypoalgesia and Hyperalgesia) and their respective neutral control conditions. Two pain modulation scores were then computed: one reflecting the hypoalgesic effect (mean pain during Hypoalgesia trials minus Neutral_HYPO_) and one reflecting the hyperalgesic effect (Hyperalgesia minus Neutral_HYPER_). These pain modulation scores were then entered into separate linear regression models using SHSS:A scores as the predictor variable. The regression effects are reported in Results, while Pearson correlations are used for visualization purposes.

### Behavioral similarity models

We used an inter-subject similarity/dissimilarity framework to characterize brain response patterns in relation to hypnotic suggestibility. This approach consists of estimating the similarity structure across all pairs of participants, and testing if subjects showing similarity in one domain (here the SHSS:A score) also express similarity in another domain (here brain responses; see *Inter-subject representational similarity analysis (IS-RSA)* below). Our main hypothesis was anchored in the Anna Karenina principle (AnnaK;^27^), which reflects an asymmetric inter-subject similarity structure where individuals located at one extreme of the behavioral continuum share strong similarity among them, while individuals at the lower end are more idiosyncratic (Fig. 2b). Specifically, AnnaK behavioral similarity was defined as the mean SHSS:A score of each dyad of participants (normalized between -1 and 1), yielding an inter-subject similarity matrix of dimension 23 x 23 with higher values reflecting more similarity at the upper end of the scale. When assessing how this similarity matrix explains the similarity structure in the neural domain, the AnnaK model can also detect the reverse pattern, i.e., higher similarity among low-suggestibility individuals. In such cases, participants with lower SHSS:A scores would exhibit more similar brain responses among them, while high scorers would exhibit more idiosyncrasies. Therefore, both positive and negative correlations are meaningful under this model, as they indicate whether a given brain region is specifically associated with stronger or weaker behavioral responses, respectively.

We also built an AnnaK similarity matrix based on behavioral pain modulation scores (e.g., Hypoalgesia - Neutral_HYPO_ scores). This model was used to validate that our findings emerging from hypnotic suggestibility as a trait shaping brain responses are also relevant for behavioral pain modulation.

Additionally, we used a second behavioral similarity model based on Euclidean distance between participants, with higher similarity scores given to dyads with similar SHSS:A scores, irrespective of the SHSS:A level. Supplementary methods detail the rationale of this model.

### Inter-subject correlations during suggestion encoding

We used intersubject correlation (ISC) analysis^58^ to assess the extent to which brain regions exhibited temporally convergent patterns of activity during auditory suggestion encoding. The fMRI BOLD signal of individuals exposed to the same dynamic stimuli comprises multiple sources of variance, and ISC isolates stimulus-locked and temporally-aligned signals across individuals^28^. Compared to the standard GLM, which assumes steady activation during a given time interval (e.g. block-design), ISC is a model-free and data-driven approach suited to capture convergent patterns of neural responses to dynamic stimuli that evolve over time, a key feature of verbal suggestions. The mean BOLD signal within 216 brain regions was extracted (200 cortical regions from the Schaeffer atlas combined with 16 subcortical regions^59,60^). ISC was then computed between all pairs of participants independently for each region. Supplementary methods describe in detail the ISC analytic procedures.

### Pain-evoked responses

We used a general linear model (GLM) to compare brain responses to painful stimulation across experimental conditions. Each participant’s design matrix included all suggestions (modelled as blocks), all shocks (modelled as discrete shock events; n = 18 per condition) and six motion parameters as nuisance regressors, plus the mean white matter estimate and mean cerebrospinal fluid. We estimated separate GLMs for each subject to produce first-level t-maps for each pain modulation contrast: Hypoalgesia vs Neutral_HYPO_ and Hyperalgesia vs Neutral_HYPER_. This contrast-based approach controls for baseline nociceptive pain processing to reveal voxelwise patterns reflecting pain modulation. These unthresholded contrast maps were used as inputs for the multivoxel spatial inter-subject representation similarity analysis.

### Inter-subject multivoxel pattern similarity during pain modulation

To assess convergent representations between subjects at the moment of pain-processing, we implemented an inter-subject multivoxel pattern similarity (IS-MVPS) approach. This method computes similarity in activation patterns across homologous brain regions for each pair of participants. Independently for each of the 216 brain regions used in the ISC analysis, we extracted the multivoxel pattern associated with each pain modulation contrast and computed the dot product between all pairs of participants. This yielded a 23 × 23 subject-by-subject similarity matrix per region, with each entry reflecting the similarity between the pain modulation patterns of two individuals. The dot product was chosen as a similarity metric because it captures both the direction and magnitude of spatial activation patterns. These IS-MVPS matrices were used in IS-RSA analyses to identify brain regions in which participants with similar behavioral scores exhibited convergent or divergent spatial representations of pain modulation.

### Inter-subject representational similarity analysis (IS-RSA)

We used representational similarity analysis (IS-RSA) to test whether inter-individual similarity in hypnotic suggestibility measured via SHSS:A scores was associated with similarity in (1) the dynamic neural patterns observed during the encoding of verbal suggestions (ISC), and (2) spatial activation patterns related to the modulation of pain (IS-MVPS). IS-RSA is specifically designed to compare pairwise similarity matrices across distinct domains^27^.

For each brain region, we computed an inter-subject similarity matrix, where each entry reflects the similarity of brain responses between two participants (Fig. 2c). These matrices were then correlated with the behavioral similarity matrix derived from suggestibility scores (SHSS:A), allowing us to test whether individuals with similar suggestibility traits exhibited more similar brain responses.

For both suggestion encoding and pain modulation IS-RSA, each neural similarity matrix was correlated to the hypnotic suggestibility similarity matrix using Spearman’s rank correlation. To assess statistical significance, we conducted two-tailed non-parametric permutation testing, shuffling the rows and columns of the behavioral matrix 10,000 times to build a null distribution. The observed correlation was then compared against this null to derive p-values. In both suggestion encoding and pain modulation IS-RSA, we tested a total of four models (2 conditions × 2 behavioral similarity models), and applied FDR correction (q < 0.05) across all 216 x 4 parcels. The resulting corrected significance thresholds were p < .0008 for suggestion encoding IS-RSA and p < .0004 for pain modulation IS-RSA.

### Cross-region IS-RSA linking suggestion encoding to pain modulation

To test how brain regions involved in suggestion encoding relate to those involved in pain modulation, we extended our IS-RSA framework to a cross-region analysis (see Fig. 2d). Specifically, we correlated ISC matrices derived from suggestion encoding (temporal IS-RSA; seed regions) with IS-MVPS matrices derived from pain modulation (spatial IS-RSA; target regions). This analysis tested whether individuals who exhibited similar temporal dynamics during suggestion encoding in a given region also showed similar spatial patterns of pain modulation in another region. Supplementary methods detail the choice of seed and target regions. Significance was assessed by applying FDR correction at q < 0.05 separately on each model x sign x condition family of tests. Corrected thresholds are specified for each cross-region set in Supplementary Table S5.

## Authors contribution

**Dylan Sutterlin-Guindon** (DSG): Conceptualization, design, implementation and sharing of Data Analysis plan (https://github.com/dylansutterlin/ISC_hypnotic_suggestions), Visualisation, Writing original draft; **Marie–Eve Picard** (MEP): Conceptualization/Revision of Analysis plan, Revising Data Analysis pipeline, Visualisation, Revision of article; **Jen-I Chen**: Conceptualization and Methodology of original study, Data acquisition and curation, Primary Data Analysis, Revision of article, Project administration, Funding acquisition; **Mathieu Landry**: Conceptualization, Revision of article, co-Supervision of DSG; **Simona Brambati**: Revision of article, Supervision of DSG; **David Ogez**: Conceptualization, Revision of article; **Mathieu Piché**: Conceptualization and Methodology of original study, Revision of article, Funding acquisition; **Pierre Rainville**: Conceptualization, Methodology, Designing/Revising Data Analysis Plan and Visualization, Revision of article, Supervision of DSG and MEP, Funding acquisition.

## Acknowledgements

D.S.G. was supported during this work by graduate studies bursaries from the Union Neurosciences et Intelligence Artificielle Québec (UNIQUE) and by Fonds de recherche du Québec—Nature et technologies. This work was supported by grants from the Canadian Institutes for Health Research (CIHR MOP-130341) and the Natural Science Research Council of Canada (NSERC RGPIN-2013-341472). We further thank Isil P. Bilgin for the valuable conceptual and analytic insights on this manuscript.

## Data availability

Open sharing of research data was not an option that participants consented to during the recruitment and primary data acquisition. Therefore, we can only share the preprocessed result outputs. Contact the corresponding author for such a request.

