## Supplementary material for "From language to pain: brain-behavior representations of hypnotic verbal suggestions for pain modulation"

### Supplementary methods

*Inter-subject correlations (ISC) during suggestion encoding*

For each of the four suggestion conditions (Hypoalgesia, $Neutral_{HYPO}$, Hyperalgesia, and $Neutral_{HYPER}$), we concatenated only the volumes acquired during a specific type of suggestion and extracted the mean BOLD time series from 200 cortical parcels[1] and 16 subcortical parcels[2] for each participant. For each region and condition, pairwise Pearson correlations were computed between all participants' time series, resulting in an ISC matrix of 23 × 23 participants with 253 unique dyads. To assess statistical significance, we used non-parametric two-sided subject-wise bootstrapping with 10,000 iterations per brain region. We tested whether the median correlation across all dyads in the ISC matrix of a given region was significantly greater than zero. Multiple comparisons were corrected using the Benjamini-Hochberg false discovery rate (FDR) procedure at $q < 0.05$ across all parcels[3]. We considered these greater-than-zero tests for each of the four conditions as a family of tests and applied the FDR correction on the number of parcels × four conditions of p-values. The resulting significance threshold was set to $p < .003$.

Furthermore, we also tested for between-condition differences in ISC using two-sided permutation testing at the subject level, again with 10,000 permutations[4]. This method accounts for the within- and between-condition correlation structure to test whether the median ISC is significantly higher in one condition than in another. The following contrasts were evaluated: Hypoalgesia vs Hyperalgesia, $Neutral_{HYPO}$ vs $Neutral_{HYPER}$, Hypoalgesia vs $Neutral_{HYPO}$, and Hyperalgesia vs $Neutral_{HYPER}$. Because ISC comparisons require an identical number of timepoints, the time series of the neutral conditions were truncated by 13 and 14 TRs to match the 100 and 101 volumes in the Hypoalgesia and Hyperalgesia conditions, respectively. FDR correction of q < 0.05 was applied across all the contrasts tested, setting significance at $p < 0.0016$.

*Euclidean similarity model of suggestibility*

As a complementary analysis to the primary Anna Karenina (AnnaK) model, we tested a Euclidean similarity model of hypnotic suggestibility. Whereas the AnnaK model captures asymmetric effects in which highly suggestible individuals exhibit more similar neural responses, the Euclidean model tests for general pairwise similarity across the suggestibility spectrum, irrespective of whether participants are high or low scorers[5]. When used in inter-subject representational similarity analysis (IS-RSA), the Euclidean model identifies non-specific contributions of hypnotic suggestibility to inter-subject similarity in brain responses.

Behavioral Euclidean similarity was computed from SHSS:A scores as 1 minus the normalized absolute difference between each participant pair. This yielded a similarity matrix ranging from 0 to 1, where larger values indicate greater similarity in hypnotic suggestibility. For visualization purposes only, similarity values were rescaled to range from −1 to 1. The resulting behavioral similarity matrix was then entered into the same IS-RSA framework used for the AnnaK analyses. Supplementary Fig. S2a illustrates how the Euclidean similarity model encodes maximum similarity between participants with adjacent scores on the behavioral scale.

*Cross-region IS-RSA for linking suggestion encoding to pain modulation*

Seed and target regions were defined based on their significant IS-RSA effects in the temporal (suggestion encoding) and spatial (pain modulation) analyses, respectively. Regions were grouped according to (i) the behavioral similarity model (AnnaK or Euclidean) and (ii) the sign of the effect (positive or negative). Cross-region IS-RSA for a given condition was computed only between seed and target regions belonging to the same model and sign, ensuring that regions showing similar inter-subject representational structure were compared (see Supplementary Table S5 for all combinations). Also note that for the Euclidean model, only positive correlation signs are valid. For each suggestion condition (Hypoalgesia, Hyperalgesia), all possible seed-target combinations within a given model x sign set were tested. For example, in the Hypoalgesia condition, AnnaK-positive seed regions (e.g., left and right anterior STG, right posterior STS) were tested against AnnaK-positive pain modulation targets (e.g., left PHG), while AnnaK-negative seed regions (e.g., aMCC, SMA, premotor regions) were tested against AnnaK-negative targets (e.g., sensorimotor areas). The complete list of seed-target pairs used for the cross-region IS-RSA is provided in Supplementary Table S5. Positive cross-region IS-RSA correlations indicate that individuals with similar ISC during suggestion encoding also exhibit similar IS-MVPS during pain modulation in another region. Statistical significance was assessed using FDR correction at $q < 0.05$, applied separately within each model × sign × condition family of tests (e.g., Hypoalgesia-AnnaK-Positive).

**Supplementary results**

*Inter-subject correlation during suggestion encoding*

To determine whether convergent brain responses during suggestion encoding were expressed across the entire suggestibility spectrum rather than preferentially among individuals with higher scores, we performed a complementary IS-RSA using the Euclidean similarity model.

During the $\text{Hypoalgesia}_{\text{CONTEXT}}$, ISC patterns in the left dorsolateral prefrontal cortex (dlPFC) were positively associated with behavioral similarity in hypnotic suggestibility ($\rho(251) = 0.21$,

FDR-corrected at $q < 0.05$; Supplementary Fig. S2b and Table S2). This finding indicates that participants with similar suggestibility scores exhibited more similar ISC in the dlPFC, regardless of whether they were high or low suggestibility scorers. However, note that the matrix structure also implies that dyads with dissimilar scores (high-low) show weaker correlation, indicating that the epochs driving the significant positive ISC in similar dyads are not the same for all the suggestibility levels.

No additional significant effects were observed for the $\text{Hyperalgesia}_{\text{CONTEXT}}$.

*IS-RSA of inter-subject multivoxel pattern similarity during pain modulation*

Using pain-modulatory contrast maps for Hypoalgesia > $\text{Neutral}_{\text{HYPO}}$, inter-subject multivoxel pattern similarity (IS-MVPS) was computed in each region, and the fit with the Euclidean similarity model was assessed using IS-RSA. The Euclidean model yielded significant IS-RSA effects in the bilateral frontal operculum/precentral gyrus (peak $\rho(251)= 0.33$), suggesting that the frontal operculum may contribute to convergent hypoalgesic responses in individuals with similar SHSS:A scores, regardless of where they fall on the suggestibility spectrum (Supplementary Fig. S2c).

For the hyperalgesic modulation, shock-evoked patterns during hyperalgesia vs $\text{Neutral}_{\text{HYPER}}$ condition were used. The Euclidean model revealed a peak in the aMCC ($\rho(251) = 0.28$), indicating that brain responses to hyperalgesia modulation in this region are influenced by inter-subject similarities in hypnotic suggestibility. Examination of the aMCC pattern in low-low and high-high suggestibility pairs of participants illustrates how similar levels of neural convergence may reflect distinct patterns across pairs of individuals on different locations of the hypnotic suggestibility spectrum (Supplementary Fig. S2d).

*Cross-region IS-RSA for linking suggestion encoding to pain modulation (Euclidean model)*

Additionally, within Euclidean-based sets of regions relevant for encoding suggestion (i.e., dlPFC), temporal alignment in the right dlPFC during $\text{Hypoalgesia}_{\text{CONTEXT}}$ suggestion correlated with spatial pattern similarity in the right precentral gyrus/frontal operculum during Hypoalgesic pain ($\rho(251) = 0.19$, $p = 0.0008$; FDR-corrected at $q < 0.05$; Supplementary Table S6). These regions had independently shown positive alignment with the Euclidean suggestibility model, indicating that brain pattern alignment scaled with inter-subject similarity in hypnotic suggestibility scores, rather than being specific to highly suggestible individuals.

*Extension of IS-RSA to behavioral changes in pain ratings*

Finally, we verified the direction of the hyperalgesia pattern in the aMCC by correlating behavioral pain hyperalgesia modulation scores to the mean activity pattern in this region. We found that individuals with the strongest activation in the aMCC showed the greatest increase in pain ratings for Hyperalgesia versus $\text{Neutral}_{\text{HYPER}}$ ($r(22) = 0.45$, $p = 0.01$; Supplementary Fig. 2e).

### Supplementary discussion

*Cross-regions associations independent of suggestibility levels*

Beyond the divergent high- and low-responder cross-regional associations from the suggestions phase to the pain modulation phase revealed by the AnnaK model, the Euclidean model identified another pathway expressed across suggestibility levels. During the encoding of $\text{Hypoalgesia}_{\text{CONTEXT}}$ suggestions, the right dlPFC showed inter-subject alignment that scaled with similarity in suggestibility scores (Fig. S2b), rather than being specific to high or low responders. This pattern is consistent with the role of the dlPFC in encoding “instructed states” and aligning cognition with top-down expectations[6,7]. Because this effect was observed across both hypoalgesic and matched neutral suggestions (context effect), it likely reflects the encoding of contextual expectations rather than modulation-specific content. However, the Euclidean structure also implies that the temporal epochs of the suggestions that drive the ISC vary with the level of suggestibility. For example, dlPFC might show more ISC among low suggestibility individuals for a particular segment of the suggestions, while this might occur during another segment in the individuals with higher suggestibility. The present study is not sufficiently powered to identify the specific epochs contributing to this difference, but future work should further examine this effect.

This shared engagement of the dlPFC during encoding was linked, via cross-region IS-RSA analysis, to similarity of hypoalgesic spatial patterns in the precentral gyrus/frontal operculum during pain. This region, part of the action-mode network involved in motor planning, posture, and arousal[8], shows that general expectancy representations formed during listening are selectively expressed in systems supporting the implementation of bodily states during pain modulation. This pathway may therefore contribute to both stronger and weaker hypoalgesic responses, depending on how expectancy representations interact with language and monitoring systems during the translation of suggestion into pain modulation.

*Hyperalgesic modulation*

In response to painful stimuli, hyperalgesic modulation produced convergent multivoxel representations in the anterior mid-cingulate cortex (aMCC) consistent with the Euclidean model, indicating that dyads with similar behavioral responses showed greater representational convergence. The aMCC is a central node in pain processing and cognitive-affective control, with activity that scales with perceived pain across experimental and clinical contexts, including hyperalgesia induced by verbal suggestions[9–11]. In our data, stronger aMCC activation and greater inter-subject representational similarity were associated with larger increases in pain ratings (Supplementary Fig. 2e), suggesting that this region contributes to the amplification of nociceptive signals during hyperalgesic suggestions compared to the control condition. Rather than reflecting a uniform response, the aMCC appears to encode hyperalgesia in a manner that varies systematically with individual differences in how suggestions are implemented. These results suggest that individual differences in expectancy and cognitive control processes contribute to hyperalgesia[12,13].

**Supplementary tables**

**Table S1.** *Inter-subject correlation comparison for verbal suggestion contexts.*

| Anatomical region | Region ID | Delta r | *p* | Coordinates (X,Y,Z) |
|---|---|---|---|---|
| **ISC difference between Hypoalgesia$_{CONTEXT}$> Hyperalgesia$_{CONTEXT}$** | | | | |
| L planum temporale | 16 | 0.2 | 0.0001 | (-53, -25, 9) |
| R planum temporale | 116 | 0.15 | 0.0001 | (52, -14, 5) |
| R post. STG | 117 | 0.13 | 0.0001 | (64, -23, 7) |
| L anterior STG | 76 | 0.12 | 0.0001 | (-56, -6, -12) |
| Right anterior STG | 187 | 0.1 | 0.0001 | (55, -6, -10) |
| Right post. STS | 189 | 0.1 | 0.0002 | (52, -31, 2) |
| L posterior STG | 78 | 0.08 | 0.0006 | (-58, -42, 7) |
| L STG | 15 | 0.08 | 0.0001 | (-51, -5, -2) |
| L paracentral lobule | 29 | 0.06 | 0.0005 | (-7, -31, 67) |
| R post. MTG | 188 | 0.05 | 0.0011 | (62, -27, -5) |
| R angular gyrus | 149 | 0.05 | 0.0007 | (60, -39, 17) |
| R parietal operculum | 119 | 0.05 | 0.0001 | (44, -27, 18) |
| L MFG | 94 | 0.04 | 0.0001 | (-24, 25, 49) |
| R ant. insula | 169 | 0.03 | 0.0011 | (34, 21, -8) |
| **ISC difference between Hyperalgesia$_{CONTEXT}$ > Hypoalgesia$_{CONTEXT}$** | | | | |
| No significant effect | | | | |

Notes: Region-wise median inter-subject correlation (ISC) values (upper triangle of the ISC matrix) were compared between conditions using 10,000 permutations to assess differences between Hypoalgesia$_{CONTEXT}$ and Hyperalgesia$_{CONTEXT}$. *Region ID* refers to the region identification number in the atlas. The atlas can be downloaded at https://github.com/dylansutterlin/ISC_hypnotic_suggestions. Coordinates of the geometric center of the cluster are in MNI space.

**Table S2.** *Hypnotic suggestibility effect on inter-subject correlations during verbal suggestion encoding.*

| Anatomical label | Region ID | Spearman rho | p | Coordinates (X,Y,Z) |
|---|---|---|---|---|
| **Hypoalgesia$_{CONTEXT}$ suggestion (AnnaK behavioral similarity model)** | | | | |
| Right anterior STG | 187 | 0.29 | 0.0001 | (55, -6, -10) |
| Right post. STS | 189 | 0.24 | 0.0001 | (52, -31, 2) |
| Left anterior STG | 76 | 0.21 | 0.0008 | (-56, -6, -12) |
| Left Premotor cortex | 28 | -0.2 | 0.0008 | (-20, -11, 68) |
| Left Somatosensory cortex | 30 | -0.21 | 0.0002 | (-19, -31, 68) |
| Right inf. Lateral Motor Cortex | 122 | -0.21 | 0.0008 | (57, -5, 30) |
| Left somatomotor | 20 | -0.22 | 0.0005 | (-56, -8, 30) |
| anterior PCC | 157 | -0.22 | 0.0003 | (11, -35, 47) |
| Right Motor Cortex | 131 | -0.23 | 0.0002 | (22, -9, 67) |
| Right pre-SMA | 158 | -0.24 | 0.0005 | (9, 4, 66) |
| SMA | 54 | -0.26 | 0.0001 | (-7, -3, 66) |
| aMCC | 52 | -0.27 | 0.0001 | (-6, 10, 41) |
| **Hypoalgesia$_{CONTEXT}$ suggestion (Euclidean behavioral similarity model)** | | | | |
| R dlPFC | 196 | 0.21 | 0.0007 | (28, 30, 43) |
| **Hyperalgesia$_{CONTEXT}$ suggestion (AnnaK behavioral similarity model)** | | | | |
| Left Supramarginal gyrus | 44 | 0.26 | 0.0001 | (-56, -40, 20) |
| **Hyperalgesia$_{CONTEXT}$ suggestion (Euclidean behavioral similarity model)** | | | | |
| No significant effect | | | | |

Notes: Region-wise correlations between hypnotic suggestibility similarity and inter-subject correlation (ISC) matrices were computed during verbal suggestion (hypoalgesia$_{CONTEXT}$ and hyperalgesia$_{CONTEXT}$) using inter-subject representational similarity analysis (IS-RSA). *AnnaK* refers to the subject-by-subject behavioral similarity model computed from pairwise means, where higher similarity is attributed to pairs of individuals with higher behavioral scores. The *Euclidean* model is the inverse of the Euclidean distance between each pair of observations. IS-RSA values (Spearman's ρ) reflect the correspondence between behavioral and brain similarity matrices, with significance assessed using 10,000 permutations. Region ID is the label for the region in the atlas. Coordinates are in MNI space.

**Table S3.** *Hypnotic suggestibility effect on inter-subject multivoxel pattern similarity during pain modulation.*

| Anatomical Labels | Region ID | Spearman rho | *p* | Coordinates (X,Y,Z) |
|---|---|---|---|---|
| **Hypoalgesia (AnnaK)** | | | | |
| Left PHG | 100 | 0.26 | 0.0001 | (-26, -32, -18) |
| Left SMA | 28 | -0.26 | 0.0001 | (-20, -11, 68) |
| **Hypoalgesia (Euclidean)** | | | | |
| Left frontal operculum | 50 | 0.33 | 0.0001 | (-51, 9, 10) |
| Right precentral gyrus | 147 | 0.21 | 0.0004 | (51, 11, 20) |
| **Hyperalgesia (AnnaK)** | | | | |
| Left OFC | 56 | 0.26 | 0.0001 | (-10, 36, -21) |
| Left OFC | 55 | 0.23 | 0.0004 | (-24, 22, -20) |
| Right motor cortex | 131 | 0.22 | 0.0006 | (22, -9, 67) |
| Left IFG | 85 | -0.23 | 0.0001 | (-46, 31, -7) |
| **Hyperalgesia (Euclidean)** | | | | |
| aMCC | 156 | 0.28 | 0.0001 | (7, 9, 41) |

Notes: Region-wise IS-RSA values (Spearman's ρ) were computed between hypnotic suggestibility similarity models (AnnaK and Euclidean) and inter-subject spatial pattern similarity during pain modulation. Conditions include Hypoalgesia (Hypoalgesia > $\text{Neutral}_{\text{HYPO}}$) and Hyperalgesia (Hyperalgesia > $\text{Neutral}_{\text{HYPER}}$) contrasts, based on GLM shock-evoked activity. Region ID is the label for the region in the atlas. Coordinates are in MNI space.

**Table S4.** List of seed and target regions included in the cross-region IS-RSA analyses.

| Model and Sign | Seed Region (Region ID) | Target Region (Region ID) |
|---|---|---|
| **Hypoalgesia$_{CONTEXT}$** | | |
| AnnaK - Positive | Left anterior STG (76) | Left PHG (100) |
| | Right anterior STG (187) | |
| | Right posterior STS (189) | |
| AnnaK - Negative | aMCC (52) | Left SMA (28) |
| | SMA (54) | |
| | Right pre-SMA (158) | |
| | Right Motor Cortex (131) | |
| | Left Somatomotor (20) | |
| | Left Somatosensory Cortex (30) | |
| | Right inferior Lateral Motor Cortex (122) | |
| | anterior PCC (157) | |
| | Left Premotor Cortex (28) | |
| Euclidean | Right dlPFC (196) | Left frontal operculum (50) |
| | | Right precentral gyrus (147) |
| **Hyperalgesia$_{CONTEXT}$** | | |
| AnnaK - Positive | Left Parietal Operculum (44) | Left OFC (55) |
| | | Left OFC (56) |
| | | Right Motor Cortex (131) |
| AnnaK - Negative | None | Left IFG (85) |
| Euclidean | None | aMCC (156) |

Notes: The seed-target sets are organized by suggestion condition (Hypoalgesia$_{CONTEXT}$, Hyperalgesia$_{CONTEXT}$), behavioral similarity model (AnnaK, Euclidean), and sign of the IS-RSA effect (positive, negative). Note that the only valid effect sign for the Euclidean model is positive. Seed regions correspond to areas showing significant inter-subject similarity during suggestion encoding (temporal IS-RSA from ISC matrices), and target regions correspond to areas showing significant inter-subject similarity during pain modulation (spatial IS-RSA from IS-MVPS matrices). Cross-region IS-RSA was computed for all seed-target combinations within each model × sign × condition set.

**Table S5.** *Significant seed-target correlations from the cross-region IS-RSA analyses.*

| Seed Region (ID) | Target Label (Region ID) | Spearman rho | *p* |
|---|---|---|---|
| **Hypoalgesia$_{CONTEXT}$ - AnnaK** | | | |
| Right anterior STG (187) | Left PHG (100) | 0.17 | 0.0033 |
| Right posterior STG (189) | Left PHG (100) | 0.14 | 0.0129 |
| **Hypoalgesia$_{CONTEXT}$ - Euclidean** | | | |
| Right dlPFC (196) | Right precentral gyrus (147) | 0.19 | 0.0008 |

Notes: Statistical significance was determined using FDR correction at $q < 0.05$, applied separately within each model × sign × condition test family (as defined in Table S4). The corresponding FDR-corrected p-value thresholds were $p < 0.01$ for the AnnaK model and $p < 0.0008$ for the Euclidean model. The region ID corresponds to the index in the Atlas used for parcellation (See Methods).

**Supplementary Table S6.** *Pain-modulatory patterns associated with AnnaK behavioral hypoalgesia*.

| Anatomical Labels | Region ID | rho | *p* | Coordinates (X,Y,Z) |
|---|---|---|---|---|
| Parahippocampal gyrus | 100 | 0.37 | 0.0001 | (-26, -32, -18) |
| Middle cingulate gyrus | 179 | 0.23 | 0.0002 | (5, 4, 30) |

Notes: IS-RSA was computed between behavioral AnnaK similarity in pain rating modulation (Hypoalgesia < Neutral$_{HYPO}$) and inter-subject spatial pattern similarity during Hypoalgesia pain modulation (GLM contrast from Hypoalgesia > Neutral$_{HYPO}$).

### Supplementary figures

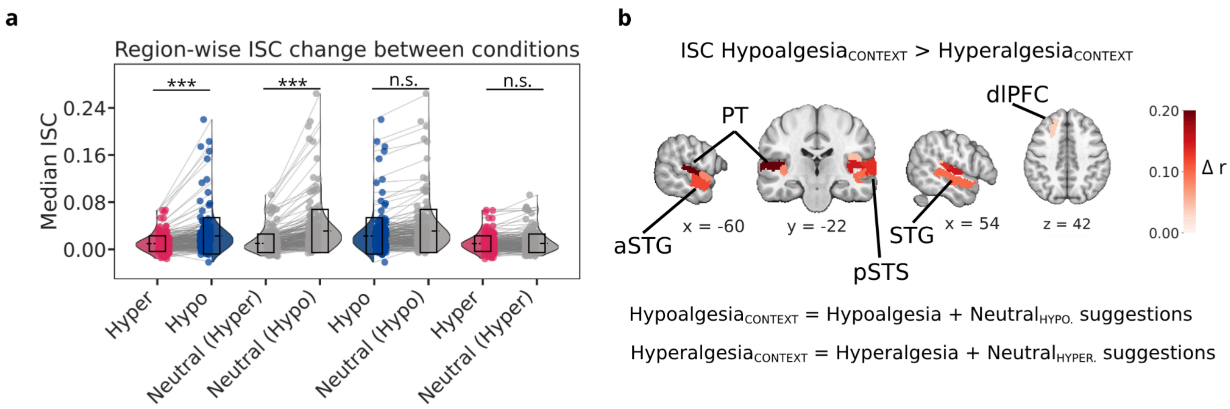


**Supplementary Figure S1.** Contextual effect of the modulatory suggestions (Hypoalgesia and Hyperalgesia) on inter-subject correlations (ISC). ISC of verbal suggestion between conditions was calculated for each of the 216 cortical parcels. **(a)** Pairwise comparisons of region-wise median ISC across conditions. Each line connects the ISC values for a single parcel across two conditions. Boxes represent the interquartile ranges, and the black dashed line indicates the median. Differences were tested using permutation-based contrast analyses[4], applied across all parcels. Significance markers reflect whether any region showed a difference after FDR correction (***p < .001; n.s., non-significant). These comparisons motivated the regrouping of conditions into $\text{Hypoalgesia}_{\text{CONTEXT}}$ (Hypoalgesia + $\text{Neutral}_{\text{HYPO}}$) and $\text{Hyperalgesia}_{\text{CONTEXT}}$ (Hyperalgesia + $\text{Neutral}_{\text{HYPER}}$). **(b)** Brain regions showing significantly higher ISC in the $\text{Hypoalgesia}_{\text{CONTEXT}}$ condition compared to $\text{Hyperalgesia}_{\text{CONTEXT}}$ (FDR-corrected at q<0.05). ISC was significantly higher for the $\text{Hypoalgesia}_{\text{CONTEXT}}$ condition (max $\Delta r = 0.20$, FDR-corrected at q<0.05), with the strongest difference in bilateral planum temporale (PT), anterior and posterior superior temporal gyrus (STG), and right posterior superior temporal sulcus (STS). Additional ISC differences were observed in the dorsolateral PFC, right anterior insula, left middle frontal gyrus, and right angular gyrus, and other regions (see supplementary Table S1).

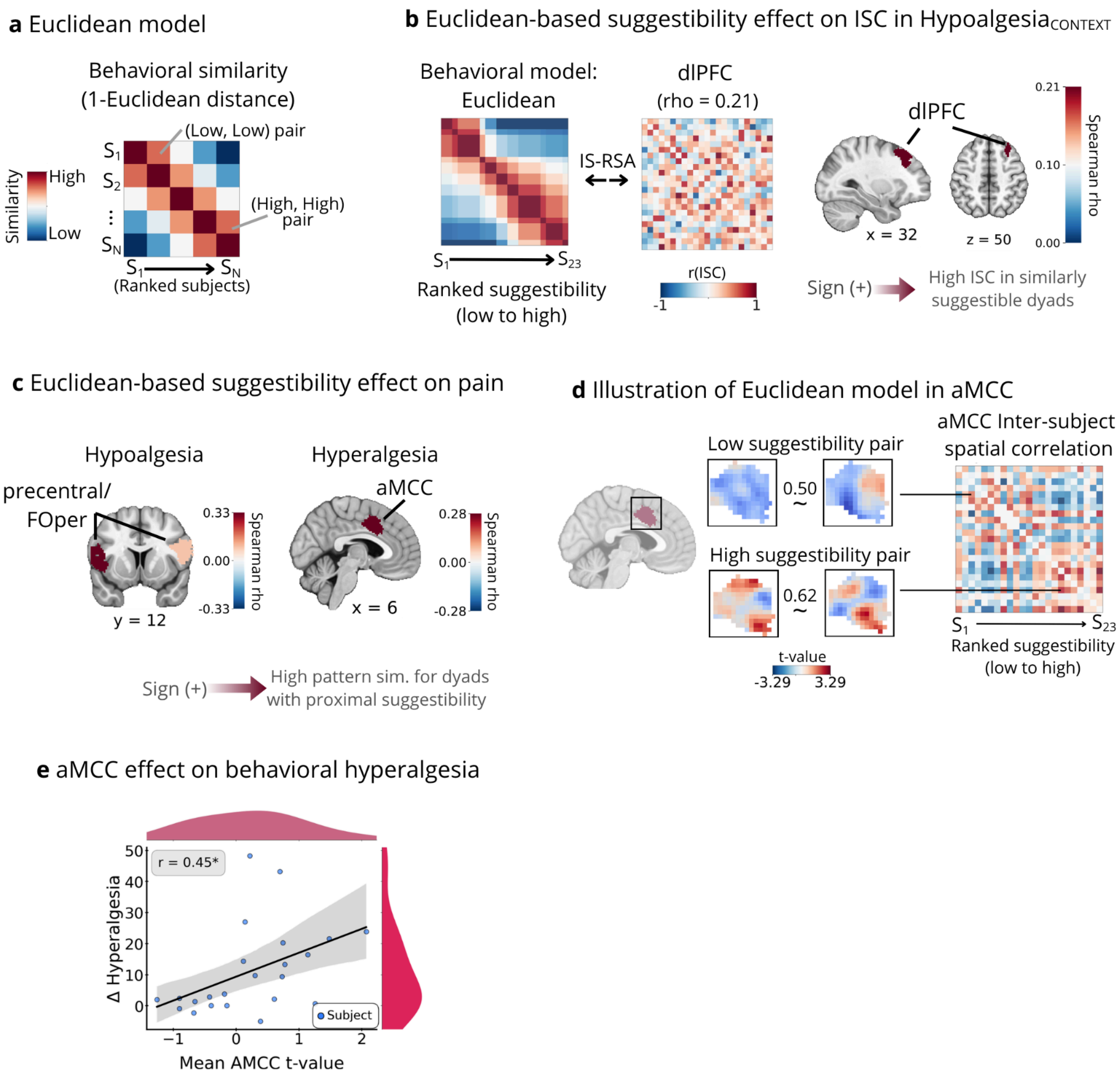


**Supplementary Figure S2.** *Euclidean-based modeling of inter-subject behavioral and neural similarity during suggestion encoding and pain modulation.* **(a)** Illustration of the Euclidean behavioral similarity model. In the Euclidean model, similarity is calculated as 1 - Euclidean distance, such that dyads with similar scores have a high similarity index, regardless of their level of suggestibility. This model is used to reveal brain regions with higher inter-subject convergence in representation for dyads with similar levels of suggestibility (e.g. high-high, moderate-moderate, and low-low) compared to dissimilar dyads (e.g. high-low). **(b)** During suggestion encoding**,** the Euclidean IS-RSA model captured non-specific effects of hypnotic suggestibility on inter-subject correlation (ISC), identifying regions where ISC was high in dyads with proximal suggestibility scores, independent of their absolute behavioral score. The dorsolateral prefrontal cortex (dlPFC) contributed non-specifically across the suggestibility spectrum to ISC scores of low- or high-suggestible dyads or participants. **(c)** During pain modulation, IS-RSA between the Euclidean suggestibility model and inter-subject multivoxel pattern similarity (IS-MVPS) of the Hypoalgesia and Hyperalgesia patterns were obtained. The Euclidean suggestibility model explained the structure of hypoalgesic pattern similarity in the bilateral precentral gyrus/frontal operculum (FOper;

left), while it explained hyperalgesia pattern similarity in the anterior mid-cingulate cortex (aMCC; right). **(d)** The aMCC is shown as a representative region where high multivoxel pattern similarity is observed both in low- and high-suggestible individuals during hyperalgesia. Examples of hyperalgesia modulation contrast maps of 2 pairs of individuals illustrate that the aMCC contributes to pain modulation across the full range of suggestibility, rather than being specific to one end of the behavioral scale. **(e)** Association between aMCC activity and behavioral hyperalgesia. Mean aMCC t-values during hyperalgesia modulation were positively associated with behavioral pain increase ($\Delta$ Hyperalgesia), indicating that greater aMCC engagement reflects stronger subjective pain amplification ($r = 0.45$).
